# Aptamer-based biosensor for food allergen determination using graphene oxide/gold nanocomposite on a paper-assisted analytical device

**DOI:** 10.1101/343368

**Authors:** Amit Tah, Jorge M. Olmos Cordero, Xuan Weng, Suresh Neethirajan

**Affiliations:** BioNano Laboratory, School of Engineering, University of Guelph, Guelph N1G 2W1, Canada

**Keywords:** Aptamers, Biosensors, Paper-based microfluidics, Graphene oxide, Food-borne allergens, Nanobiosensing platform

## Abstract

The detection of allergens in food are currently conducted by techniques that are time-consuming and complicated which can deter consistent sampling for allergens, which could potentially cause an anaphylactic shock in the consumer by cross-contamination. The need for a technique that is rapid, on-site, cost-effective, disposable, highly sensitive and accurate to identify these molecules urges the development of a point-of-care device. The aim of this work is to develop a microfluidic paper-assisted analytical device (μPAD) using hydrophobic channels, set by a wax printer on filter paper, and functionalized gold nanoparticles (AuNP) to help identify the allergens arachin (Ara h 1) for peanuts, β-lactoglobulin (β LG) for milk, and tropomyosin (Pen a 1) for shrimp and other shellfish presence by a colorimetric test. Synthesized AuNP were conjugated with biotinylated aptamers, using the biotin-streptavidin interaction, to make the specific detection of target allergens. Functionalized AuNP are incubated with the sample and are absorbed by graphene oxide (GO), creating GO-AuNP complexes, if the aptamers have not become structured due to conjugation with allergenic proteins. The μPAD device is used to filter the resultant mixture which provides superior sensitivity to detect the allergens present down to the nanogram range (allergens were measured from 25 nM - 1000 nM with a LOD of 7.8 nM, 12.4 nM and 6.2 nM for peanut, milk and shrimp allergens respectively), in contrast to the microgram range of commonly used enzymatic immunoassays. The simple color indicator, varying from clear to pink in the presences of allergens allows the readout to be utilized without the need for highly specific equipment or training. Alternatively, the results can be quantified by taking a picture and measuring the color. This presented μPAD can provide results in real time and has the potential to become a rapid, low-cost, and accurate portable point-of-care device to avoid cross-reactivity of food-borne allergens.

## 1. Introduction

Food allergies are caused when the immunological system of a person overreacts adversely to consumed proteins found in the meal denoted as allergens (AAFA, 2017). The symptoms an individual experience from an allergic reaction vary from one to another and can range from mild to severe. Unfortunately, the causes for allergies are not yet completely understood but the mechanisms are reported and thus allow for precautionary measurements to be taken (Smith, 2013). There is no current cure to allergies thus the allergic reactions are prevented by avoiding contact with the food, controlling the symptoms (taking antihistamines, epinephrine) and allergy injection therapy (CDC).

FDA and other governmental bodies identify 8 major allergens that affect American society which include milk, eggs, fish, crustacean shellfish, tree nuts, wheat and soy (Center for Food Safety and Applied Nutrition, 2017). However, major allergens can vary depending in the population and the region of the world thus they are the ones considered to evoke a reaction in more than 50% of allergic patients (Kallós et al., 1978). In the United States alone 15 million individuals are sensitive to the various food borne allergens (FARE, 2016). To prevent accidental reaction sensitive individuals must rely on manufactures, governmental regulation and imposing strict diet on themselves to lower the chance of a response due to allergenic material (World Allergy Organization, 2011). In this study, peanut, milk and shellfish were selected due to being the most common food allergens in children in the U.S. with an incidence of 25.5%, 21.1% and 17.2%, respectively. It is important to mention that out of the children some had reported having a history of a severe reaction and 30.4% were allergic to more than one allergen. Ara h 1 was chosen as the peanut allergen due to its abundant 23 independent IgE-binding epitopes (Zhuang and Dreskin, 2013). Choosing a milk allergen is not as simple given that most of cow’s milk proteins have potential allergenic properties thus β-lactoglobulin was chosen for its abundance (Natale et al., 2004). Tropomyosin was chosen for shrimp detection given that it is very frequent and it provides a simultaneous test for contamination with dust mites and cockroaches for people who are also sensitive to arthropods (Pascal et al., 2015). With so many sources and individuals varying sensitivity, reactionary based methods like EpiPens, are not the most ideal mechanism for helping those afflicted. To meet the need for proactive measures which can be utilized directly by consumers on-site field testing allergen detection tool is required.

The advent of paper-based microfluidic, which utilizes the natural capillary action of cellulosic substrates to perform rapid diagnostic test, may change this and help bring the Point-of-Care testing (POCT) to the masses (Martinez et al., 2010). This new type of diagnostic system provides a methodology which can help take microfluidics from the lab to the consumers. The advantages of paper-based microfluidics includes the low cost, ease of manufacturing, the possibility of simple multiplexing, small sample/reagent consumption and capability for development of a three dimensionally structured diagnostic tool (Martinez et al., 2007). These advantages allow paper based devices to deliver on the promises that original lab-on-a-chip microfluidic put forward, the development of POCT for the entire world.

For this device, the main sensing mechanism incorporates the use of aptamers to accurately identify the allergenic proteins. Aptamers are short single-stranded DNA or RNA oligonucleotides that have the ability of binding to specific molecules showing an activity similar to antibodies. The structure of aptamers contains complementary base pairs that allow a stable secondary structure to form a rigid functional structure to bind with their target molecule. There have been aptamers developed to identify metal ions, small organic molecules, peptides, proteins, viruses, bacteria, and whole cells (Keefe et al., 2010). Aptamers, thus prove to have advantages on detection over antibodies by being smaller in size (6-30 kDa or 20-100 nt), having a high affinity independent from the number of epitopes in the target molecule, allowing to identify single point mutation and isomers, detecting a wider range of molecules, rapid production, low variation between batches, low risk of contamination, long stability, having the ability to be modified, and low to none immunogenicity (BasePair Biotechnologies, 2017; Zhou and Rossi, 2016). They can be used in biosensors with a higher density than antibodies and have been proven to be reusable without changes in specificity or sensitivity (Lakhin et al., 2013).

In our study, aptamers are conjugated to gold nanoparticles (AuNP), we then utilize specific single stranded DNA (ssDNA) absorption properties of graphene oxide (GO) with a paper device to create a simple colorimetric sensor for food allergen detection. These two materials, AuNP and GO, have been highly exploited for the development of POCT in recent years thanks to their fantastic properties and interactions. AuNP have very useful optical and electrical properties, their strong and specific Surface Plasmon Resonance absorption and extremely high extinction coefficients allows them to be ideal reporters for target analytes with low cost equipment or with the naked eye (Alves et al., 2016). GO is also a wonder material in recent years thanks to its ability to quench fluorophore-labeled bio-recognition molecules, such as labelled antibodies, which when in the presence of their target reverse the quenching and re-release their fluorescent signal due to F_rster resonance energy transfer (FRET) (Lu et al., 2015). It also has been shown to interact with a variety of biological molecules such as amino acids, peptides, proteins and most importantly for this application ssDNA (Li et al., 2014, 2012). It has been shown that ssDNA has a special adsorption interaction with GO, forming a complex in which the functionalized AuNP have a bridging effect between different layers of graphene oxide through pi-pi stacking thanks to the ssDNA aptamers bound to them. In contrast, double stranded DNA or highly structure molecules of DNA do not show the formation of pi-pi interaction with GO, as their complex-rigid structure prevents the pi-pi interaction from occurring with GO (Li et al., 2014). By tagging ssDNA aptamers, which are designed to detect target allergens, to AuNP it is possible to create a colorimetric biosensor which can produce an output visible to the naked eye. Combining this with a simple paper based device, it allows an easy and fast quantifiable assessment of allergen concentration. The mechanism of the presented biosensor and resultant reaction for allergenic proteins in food is shown in Fig. 1.

**Fig. 1.**
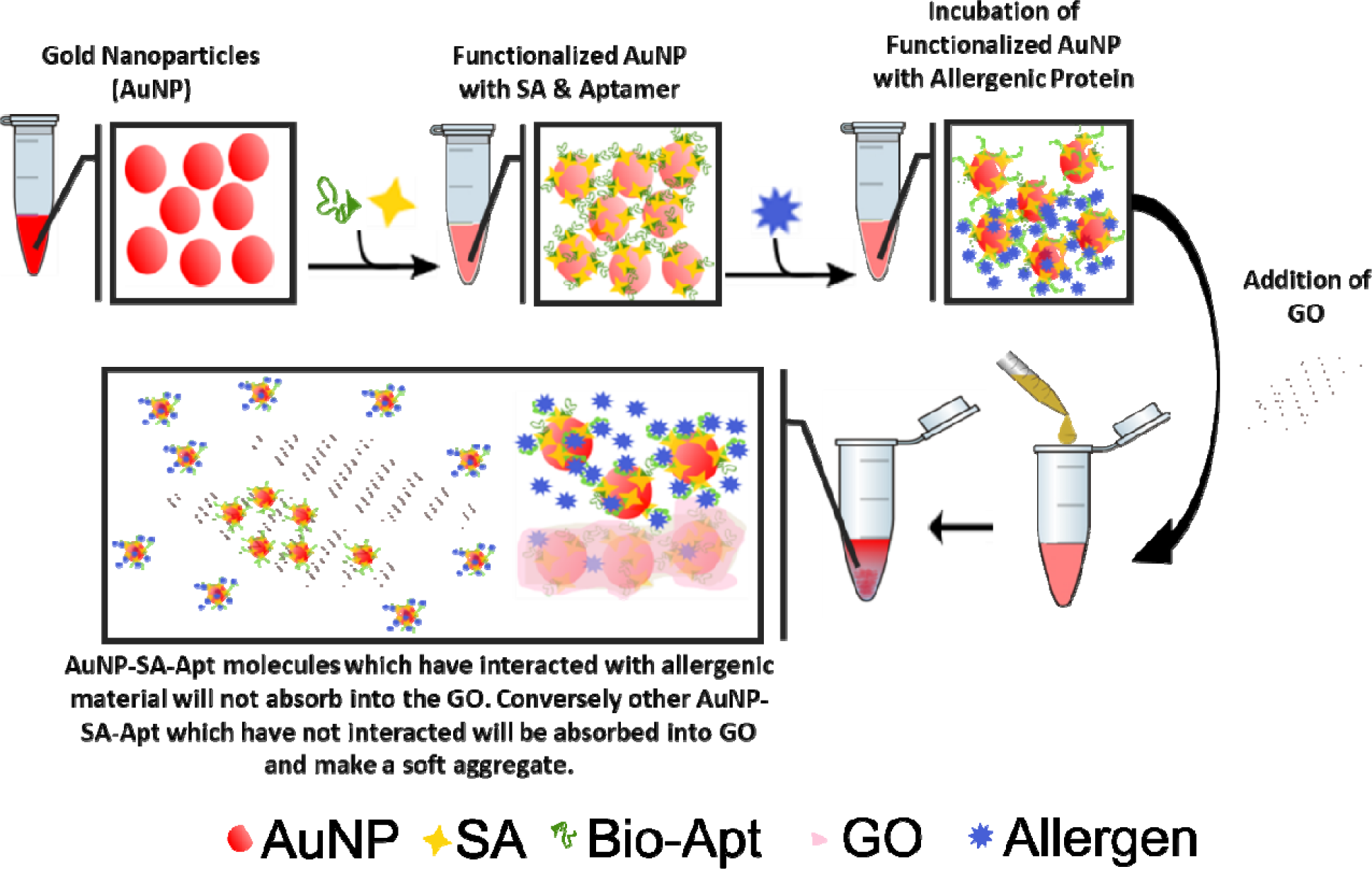
Schematic of the proposed mechanism for colorimetric detection of allergens with the use of AuNP, Aptamers and GO. When in the presence of allergenic proteins the AuNP-SA-Apt molecules are prevented from interfacing with the GO when added, if some AuNP-SA-Apt molecules have not

## 2. Materials and methods

### 2.1. Materials

Tetracholoroauric(III) acid (HAuCl_4_ • 3H_2_O), trisodium citrate (Na_3_C_6_H_5_O_7_ • 2H_2_O), streptavidin, β-lactoglobulin (βLG), phosphate buffer saline (1x PBS; 0.01 M phosphate buffer, 0.0027 M potassium chloride, 0.137 M sodium chloride, pH 7.4), graphene oxide, bovine serum albumin (BSA), sodium chloride (NaCl), magnesium chloride (MgCl_2_), tris(hydroxymethyl)aminomethane (Tris, C_4_H_11_O_3_N), ethylenediaminetetraacetic acid (EDTA, C10H16O8N2), mixed cellulose ester filters (MCE, 0.45 μm), boric acid (H3BO3), sodium hydroxide (NaOH), Tween 20 and Whatman chromatography paper (cellulose, 15 cm x 100 m) were purchased from Sigma-Aldrich Canada (Burlington, ON, Canada). Arachin (Ara h 1) and tropomyosin (Pen a 1) were purchased from Indoor Biotechnologies. Aptamers were synthesized by Integrated DNA Technologies (Coralville, Iowa, USA). The sequence of the aptamers is listed in the Table 1.

**Table 1.**
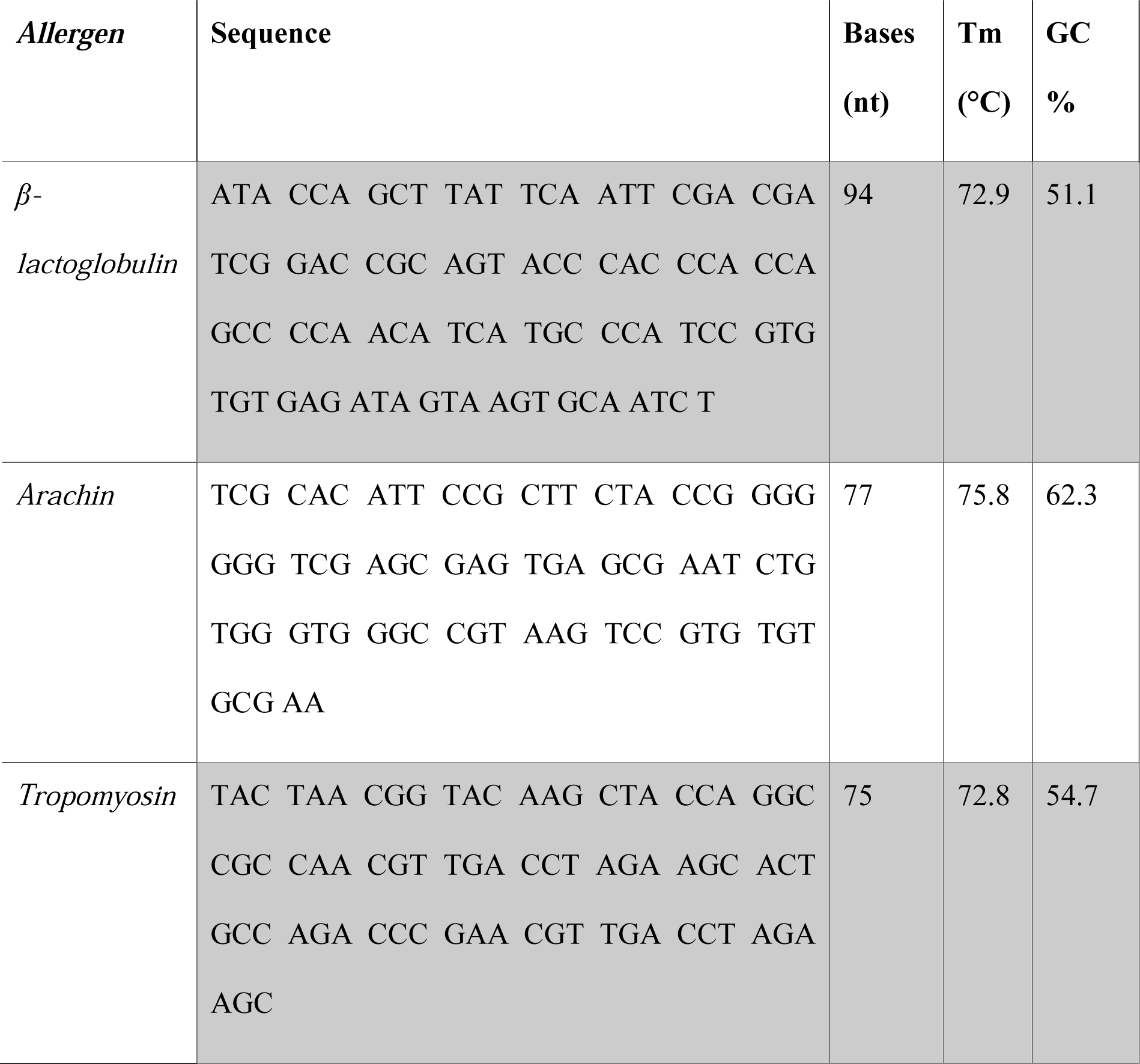
Aptamer Sequence for detection of each specific allergen

### 2.2. Equipment

TEM imaging (Tecnai G2 F20) was employed to obtain the micrographs. For the spectral scanning, Cytation 5 (BioTek), and Cary 100 UV-Vis (Agilent) were used. Dynamic light scattering measurements were conducted using Brookhaven 90plus Particle Size Analyzer (Brookhaven Instruments Corporation). The μPAD,s hydrophobic channels were constructed with the wax printer Colorqube 8580 (Xerox) and the hot plate (Sci Logex MS7-H550-S). The samples were centrifuged on Sorvall ST 40 R (ThermoFisher Scientific). The aptamers were heated in Precision Water Baths 280 (ThermoFischer Scientific). Incubations took place in the Incushaker mini (Southwest Science). Milli-Q water was used in all the experiments (0.055 μS/cm).

### 2.3. Gold nanoparticles synthesis

Gold nanoparticles were prepared using Turkevich’s method (Turkevich et al., 1951). A 100 mL of HAuCl_4_ • 3H_2_O (1 mM) were boiled while stirring in a hot plate with a magnetic bar. Then 10 mL of Na_3_C_6_H_5_O_7_ • 2H_2_O 38.8 mM were added. The reaction was left to boil for 10 minutes until the color changed from yellow to black to deep red wine. AuNP were left at room temperature for cooling down, prior to filtration with the MCE membranes. It was required for the nanoparticles to be covered from light and kept at 4°C to ensure their stability. The nanoparticles have a SPR peak at 521 nm indicating a 15-nm diameter size which is in correspondence with literature (Haiss et al., 2007). Theoretical calculation state that the synthesised particles are expected to be dispersed as a 1.1 nM concentration, which was calculated with an estimated extinction coefficient of 2.18X 10^8^ (Haiss et al., 2007).

### 2.4. Streptavidin conjugation (AuNP-SA)

Thawed 1 mg/mL streptavidin was diluted to 50 μg/mL in 400 μL of borate buffer (0.1 M, pH 7.4). Conjugation with 600 μL of AuNP was performed at 4°C in moderate shaking for 30 minutes. To remove the unbound streptavidin, AuNP-SA were centrifuged for 40 minutes at 4500 rpm and 4°C. The supernatant was discarded and the pellet was resuspended in 1x PBS. A second wash was performed in the same conditions. The final resuspension was made in 100 μL of PBS (Lim et al., 2012).

### 2.5. Aptamer functionalization and conjugation (AuNP-SA-M/P/S)

Lyophilized aptamers were centrifuged for a pulse and then resuspended in TE buffer (10 mM Tris, 0.1 mM EDTA, pH 7.5) as per the protocols provided by Integrated DNA Technologies to 100 μ The aptamers were incubated at room temperature for 30 minutes and vortexed. Afterwards, the aptamers were pulsed at 10,000 g for 2 seconds and aliquoted to be preserved at −20°C. Prior to be used, aptamers were diluted to 50 nM in 100 μL of folding buffer (1 mM MgCl_2_, 1x PBS) and denatured at 90°C for 5 minutes. They were let to cool down at room temperature for 15 minutes. Streptavidin coated gold nanoparticles were mixed with aptamers at room temperature for 30 minutes under gentle mixing. Excess aptamers were removed by centrifugation for 15 minutes at 6000 rpm and 4°C. The pellet was resuspended in 1x PBS, repeating the washes thrice (Weng and Neethirajan, 2016). Conjugated AuNP-SA-M/P/S were left for 16 hours to age at 4°C.

### 2.6. Graphene oxide absorption of functionalized AuNP (AuNP-SA-M/P/S + GO)

One of the critical mechanisms is the absorption of ssDNA with the usage of GO after the functionalized AuNP are incubated with a sample. Functionalized AuNP which have not had interaction of the aptamers with allergenic proteins are absorbed and form soft complexes. A total of 100 μL of nanoparticles were mixed with 0.02 mg/mL graphene oxide of which is extensively sonicated for 30 minutes before usage.

### 2.7. Paper-based analytical device

For this device, paper tests were made on Whatman chromatography paper with designs from Inkscape (0.92, The Inkscape Project, open-source program) using Colorqube wax printer. The paper coated with the device was heated in the hot plate at 170°C for 2 minutes to create the hydrophobic channels using a glass slide to cover the paper system (Lee and Gomez, 2017). The paper was cooled down prior to loading of the supernatant of the AuNP-Apt + GO complex and solution. The folded paper system is now ready to direct and channel small volumes of the supernatant to filter and display the testing results.

### 2.8. Characterization and validation

The physical diameter of the nanoparticles was determined by analyzing TEM micrographs with ImageJ (Schneider et al., 2012). 20 μL of diluted samples were placed on copper grids and let sit overnight to be absorbed before imaging with the electron microscope. The analysis of the images included applying a FFT Bandpass Filter making large structures down to 20 pixels and small structures up to 5 pixels. Then the image was adjusted using the threshold tool and the particles’ area was analyzed with a circularity above 0.3.

Dynamic Light Scattering (DLS) measurement was taken by adding 200 μL of filtered samples in deionized water in the cuvettes and diluting as the count rate reach an approximate value of 400 kcps.

Absorbance was measured using Cytation 5 or Cary 100 using a 96-well plate or cuvettes. A 10Χ dilution factor was used for all scans unless otherwise noted, all necessary graphs will show the dilution correct values. Every step of the conjugation was verified with data obtained in Cytation 5 and data was further processed as necessary.

## 3. Results and discussion

### 3.1. Characterization of functionalized AuNP

To demonstrate that the biomolecules were being attached to AuNP, TEM images and DLS measurements were acquired. For characterization of the mechanism, the data represented below is with the conjugation of Beta-Lacto Globulin specific aptamer. Each conjugation step was expected to occur on the surface of the AuNP without causing seeding effects, or a change in morphology. Nanoparticles preserved their spherical morphology and the average diameter (~14 nm) from AuNP upon conjugation with SA and biotinylated aptamers, respectively (Fig. 2). The hydrodynamic radii were obtained with the use of DLS giving values of 24.2 nm, 45.7 nm, and nm for AuNP, AuNP-SA, and AuNP-SA-M respectively. Unlike the TEM which is unable to visually image the conjugated proteins and ssDNA, the DLS analysis shows an increase in the hydrodynamic radii from each functionalization. Zeta analysis values were also obtained giving values of −72.41, −38.74 and −46.54 for AuNP, AuNP-SA, and AuNP-SA-M/P/S, respectively. The increase in absolute potential seen when AuNP-SA are further functionalized with the aptamer shows a change in the surface morphology with a more negative molecule, which further attributes to the biotinylated aptamer binding with the SA (D’Agata et al., 2017; Weng and Neethirajan, 2016). From Fig. 2F, spectral scanning shows SPR peaks of conjugated 14 nm AuNP from 520 nm to 531 nm when covered with SA and to 536 nm when aged aptamers were absorbed onto SA. The adsorption of protein on to the surface causes a change in the surface morphology and therefore effects the scattering of the particles. The well-known optical characteristics, which were first systematically explained by Mie (1908), describe how the localized environment affect these properties. As gold has a strong Surface Plasmon Resonance, the shift of the peak can be directly correlated to a change in the localized refractive index. Therefore, a shift in the spectral peak of gold can be a correspondence to a new molecule which interferes with the natural gold Surface Plasmon Resonance peak. The introduction of the protein SA and its electrostatic adoption onto the surface causes this change in the local refractive index, which in turn causes the shift of the spectral absorbance (Pollitt et al., 2015). The second shift is caused by the introduction of the biotinylated aptamers, which directly bind to the SA, leading to a change in the SA localized refractive index (D’Agata et al., 2017; Pollitt et al., 2015). Inserts on Fig. 2A, 2B & 2C demonstrates the diameter distribution of the functionalized AuNP from the processed micrographs to ensure that the peak shift to 536 nm does not correspond to flocculation but rather to surface biomolecule addition. Using a combination of DLS analysis, spectral scanning and TEM studies, a determination that surface modification of the nanoparticles due to the conjugation of the bio-molecules (SA and aptamers) has been successful, creating stable and fully functional nanoparticles for the detection of allergens.

**Fig. 2.**
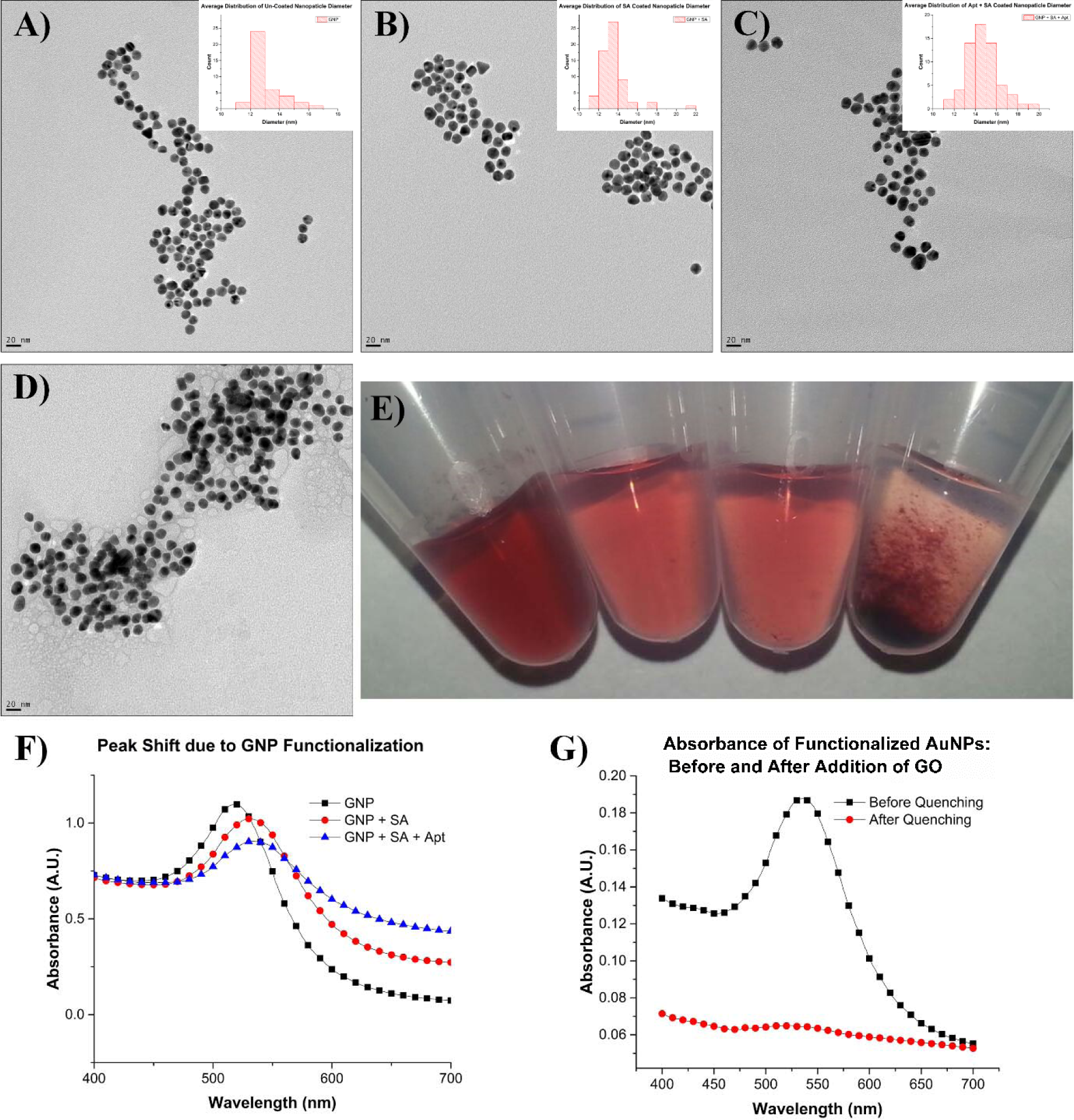
TEM micrographs of A) AuNP, B) AuNP-SA, C) AuNP-SA-Milk Apt, and D) AuNP-SA-Milk Apt +GO nanosheets with histogram of average diameter found to right. E) Image of AuNP, AuNP-SA-ssDNA, AuNP-SA-ssDNA + GO + 1000 nM allergen spiked sample in solution and AuNP-SA-ssDNA + GO with no allergen spike F) Spectral shift caused by the conjugation of SA and aptamer compared to bare nanoparticles. G) Spectra of functionalized AuNP before and after the addition of GO.

Fig. 2D and 2G show and explain the effect of nanoparticles being bound to sheets of graphene oxide, creating bridges between sheets, resulting in the complexes seen in Fig. 2D. As explained previously, the pi-pi stacking interaction between the different layers of graphene oxide due to the functionalized AuNP only readily occurs when the aptamers are unstructured (Park et al., 2014; Wang et al., 2011; Wu et al., 2011). As no allergenic proteins are present, most nanoparticles have been removed from solution, as seen from the spectrographs of the supernatant in Fig. 2G. The background layer of graphene oxide and the implanted nanoparticles in Fig. 2D further support this assessment of interaction between the nanoparticles.

### 3.2. Optimization

The proposed mechanism has several critical steps that can be changed to improve the reliability, reproducibility and maximize the signal that is produced. The first study was assessment of different nanoparticles synthesis methods to create various mean diameters. Turkevich’s method (Kimling et al., 2006) was finally chosen due to yielding consistent sizes and bigger particles than Martin’s method (Low and Bansal., 2010) thus proving to have enough surface area to chemisorb SA. Based on previous studies of SA, it was estimated that between 20 nm^2^ ~ 40 nm^2^ of surface area is required for electrostatic adsorption of SA (Bayer et al., 1990). As stated above, the simple and consistent Turkevich method provided a nanoparticle of consistent size with ample surface area for adsorption of SA molecules. Based on TEM, there was an estimated average available surface area of approximately 620 nm^2^. Next, choosing the correct working buffer became an important condition when conjugating our functionalized AuNP; PBS was only used after ensuring AuNP had the critical concentration of SA thus making the first conjugation in borate buffer non-saline mandatory (Geneviève et al., 2007). This was because borate buffer (0.1 M pH 7.4, 0.01 M pH 7.4 and 0.1 M pH 6.4) did not have any adverse effect on AuNP; however, PB 1* pH 7.4 and SC (sodium citrate) 1* pH 7.0 cause instability and eventual flocculation of the nanoparticles. This prevented any interference from occurring when the first functionalization with SA occurred. The critical concentration of SA was determined with the use of a salting flocculation test (Geneviève et al., 2007) and the stability verified by the SPR peak shift (Lim et al., 2012). The monolayer production test and optimization data can be seen in Fig. 3A and 3B, resulting in the optimum concentration to provide a monolayer of SA for AuNP to be 50 μg/mL of SA. To optimize the aptamer concentration, we first theoretically calculated the minimum amount required for total coverage of the SA cover AuNP by the method introduced in previous studies (Pollitt et al., 2015; D’Agata et al., 2017; Sechi et al., 2013).

**Fig. 3.**
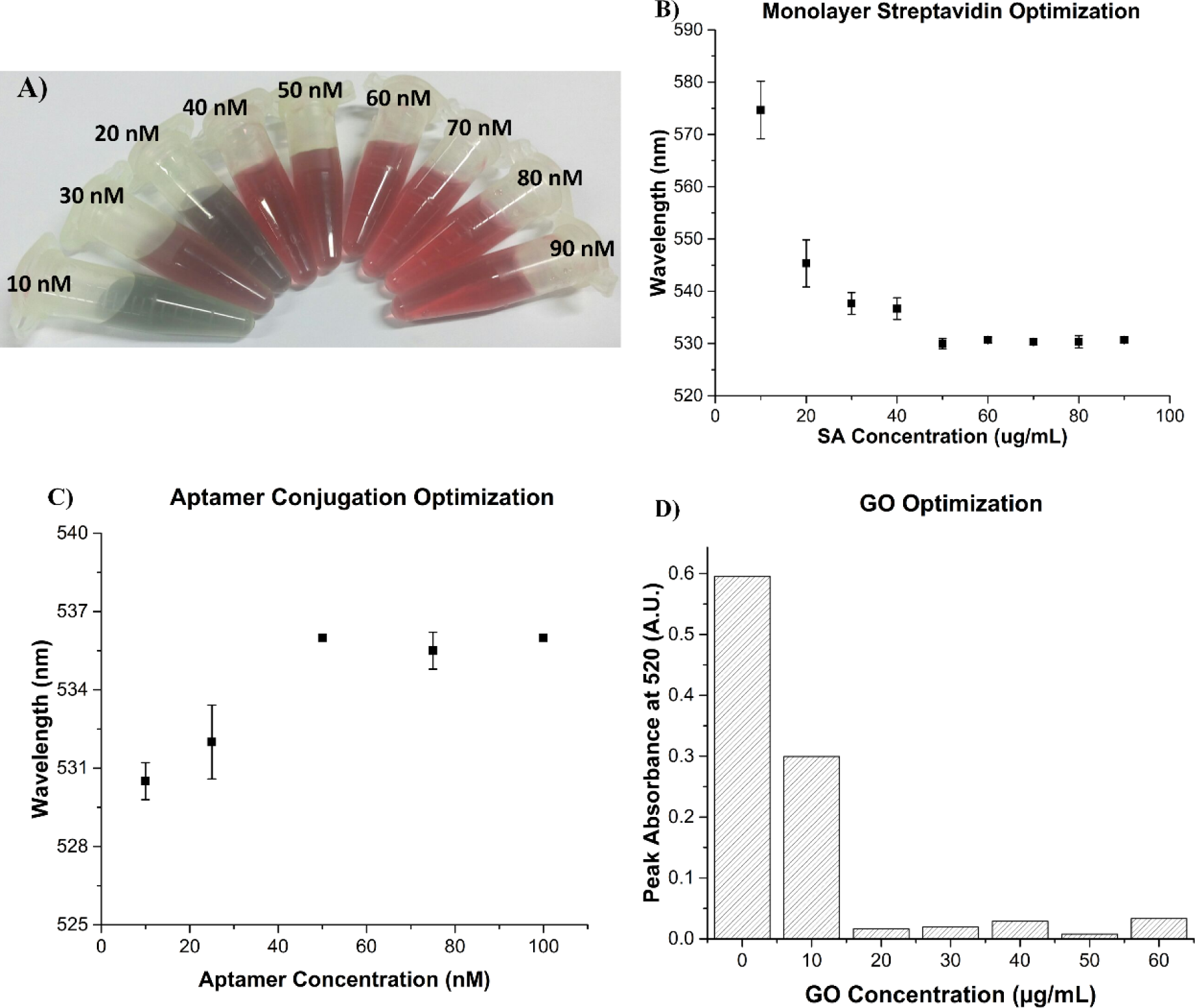
A) SA conjugation from 10 μg/mL to 90 μg/mL after NaCl addition. B) Spectra shift after SA conjugation. C) Spectral shift after aptamer conjugation using different concentrations. D) GO optimization conjugation using the peak absorbance at 520 nm with 45-minute wait after introduction of GO.

From these studies, an estimated 23 molecules of SA are available on each AuNP, which correlates to approximately 46 available biotin binding sites for biotinylated aptamers. While it is known that SA has 4 possible biotin binding sites, it was estimated that 2 would be incapable of binding with biotin. This assumption is based on the work conducted by D’Agata et al., which listed this effect is related to steric obstruction of binding sites caused by the most likely orientation of absorption for SA onto the surface of the NP (D’Agata et al., 2017). With our known initial concentration of AuNP, an estimate of 38 nM solution of biotinylated aptamers is needed for the total coverage. To prove this estimation, a similar procedure to the SA optimization was used to determine the optimum concentration of aptamers (Lim et al., 2012), and as seen in Fig 3C, 50 nM of biotinylated aptamers yielded to a stable shift in the spectra of 536 nm. Although not an exact estimation we can see that the minimum possible values of aptamer needed is between 38 nM and 50 nM for complete and stable functionalization.

For the mechanism to properly detect their target allergen, GO concentration was also optimized by varying the concentration from 10 μg/mL to 90 μg/mL and getting the spectral scan from the supernatant after 45-minute wait and a pulse centrifugation at 6000 rpm for 10 seconds. This optimization was done in accordance with minimizing the amount of GO needed as it is also capable of interacting with SA, which could cause a similar reduction of fully functionalized nanoparticles (with aptamers) found in the supernatant if given enough time (Li et al., 2012). Therefore, to limit the loss of the colorimetric signal we optimized to use as little GO as needed. This was done because ssDNA such as aptamers, when unstructured, are much more sensitive and capable of creating the necessary pi-pi interaction required in comparison to structured proteins like SA (Li et al., 2014, 2012). However, interaction is possible so the decision to minimize the amount of GO used for both decreasing the cost and increasing specificity of the output was prioritized. The minimum concentration that allows the mechanism to fully absorb unconjugated functionalized AuNP within the 45-min reaction time is 20 μg/mL of previously extensively sonicated GO (30 minute of prior sonication) (Fig. 3D).

### 3.3. Biosensor validation, selectivity, and sensitivity

To verify that the mechanism works as expected in solution, trials were first conducted to determine consistent response to given concentrations of allergens. For these tests, a spiked sample of protein allergen was introduced to the fully functionalized AuNP at different concentrations. Then GO was added at the previously optimized amount and 45 minutes of time was given to allow formation of the aggregates with the GO and AuNP. When the aptamer interacts with its specified protein, its structure changes from one which can interact with GO and create pi-pi stacking in to a rigid structure which weakly produces these bonds (Li et al., 2014). Through this mechanism of interaction, and the ability to prevent interaction we can selectively remove functionalized AuNP which have not conjugated with the specific allergenic protein from the solution with GO. This will create a colorimetric signal which is proportional to the amount of allergenic protein found in the food sample solution. This hypothesis was proven as shown in Fig. 4B where a standard curve for each aptamer is shown. The logarithmic model fit applied to peanut, milk and shrimp allergen response were .99474, .94134 and .98149 respectively.

**Fig. 4.**
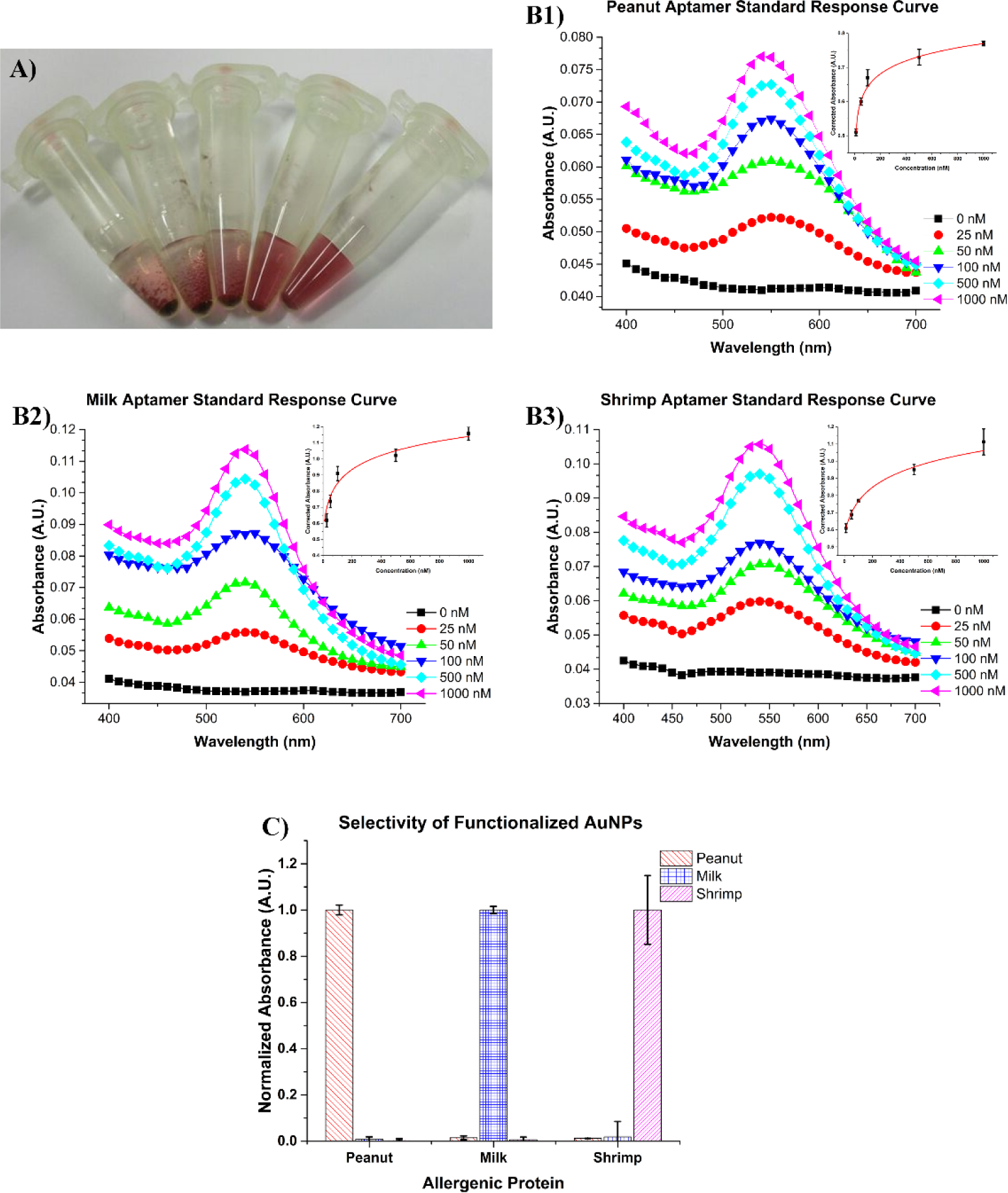
A) Functionalized AuNP reacted with (from left to right) 25 nM, 50 nM, 100 nM, 500 nM and 1000 nM of allergenic protein and pulse centrifuged. B1, 2&3) Spectral signal from the supernatant due to the reaction and GO absorption of the unstructured aptamer on AuNP from various allergen concentrations of B1) Peanut, B2) Milk and B3) Shrimp, bottom to top 25 nM, 50 nM, 100 nM, 500 nM and 1000 nM. C) Selectivity study of the functionalized AuNP system to the other allergenic proteins.

Using various concentrations of allergenic proteins, we can see a clear relationship between the allergen concentration and amount of AuNP left in solution. In testing, we measured allergen concentrations down to 25 nM. The calculated lower limit of detection (LoD) for each allergen detection systems were calculated with a 3σ ratio, calculated to be 7.8 nM, 12.4 nM and 6.3 nM for peanut, milk and shrimp systems respectively (Thomsen et al, 2003). As described by Thomsen et al. (2003), limits of detection were specifically calculated by first calculating the sensitivity (S) of system which was the change in concentration over the mean change of signal within the linear region. Then the standard deviation of the blank sample trials (S_bl_) was calculated and used in the equation C_L_ = k*(S_bl_)*(AConcentration / AIntensity) (Thomsen et al. (2003)) with a 3σ noise ratio (k). The FDA threshold level in food cross-contaminates are 0.25 mg, 0.36 mg and 0.13 mg of peanut, milk and shrimp allergenic proteins respectively (Center for Food Safety and Applied Nutrition, 2016). The developed system can detect concentrations well under this required limit. This is a significant benefit as sensitive individuals may have different levels of sensitivities to these allergens. Although for the lower limit a spectrometer is required, for industrial application in the food industry, our developed tool provides a fast-quantitative alternative for current conventional methods like ELISA; which also require bench-top devices and significantly more reagent volumes and steps. Most ELISA kits offered by major suppliers like NEOGEN or r-Biopharms require bench top equipment like plate readers, a significant number of reagents and take 30 minutes or longer (NEOGEN, 2017a, 2017b, 2017c; r-Biopharm, 2017a, 2017b, 2017c). These kits are considered the gold standard for industry; however, most are only able to detect within the microgram range in comparison to the mechanisms’ ability to function at nanogram levels. Another significant benefit of this system is the simple processes required to obtain a measurement, in comparison to the multiple reagent ELISA tests that these systems utilize for detection of allergenic material, the proposed mechanism only requires operators to add GO after the sample has been incubated with the pre-functionalized AuNP. The specific affinity for their target proteins is seen in Fig. 4C, where a selectivity study shows the output of the system which are spiked with 1000 nM of each protein. Although more comprehensive cross reactivity studies are necessary, there is limited cross reactivity between these aptamers. This makes them ideal for their application as an onsite biosensor as they are both highly sensitive and selective, with results that directly correspond to the amount of allergen present.

### 3.4. Microfluidics paper-based analytical device

Paper based microfluidics offer a variety of benefits which can be utilized to make the proposed mechanism much more available outside the laboratory environment. The main goal of the proposed bio-sensing system is to increase the options for a sensitive individual or food manufacturer to quickly and efficiently detect these proteins. To accomplish this, we utilized simple cellulose based paper to both help separate the created complexes from solution and increase the readability of the result. Using paper to filter the sample matrix is one of its oldest known usage in science (Yetisen et al., 2013), and here we can use it for this specific purpose. We selected Whatman 1 cellulose based paper as it does not degrade over time, its availability and low cost (GE Healthcare Life Sciences, 2017; Mahato et al., 2017). To first verify if it is possible to filter the developed complexes with Whatman paper (pore diameter of 11 μm) test samples were first incubated with varying quantities of allergenic protein and then allowed to interact with GO and filtered through the paper. The supernatant derived from this test can be seen in Fig. 5A, and shows the ability of the paper to remove the developed complexes from solution, while allowing passage way for the un-absorbed AuNP. As we know that Whatman 1 filter paper can selectively remove the complex from solution we designed a paper device which utilizes a simple 3-fold design, as seen in Fig. 5B, to filter and present the output of the results. We added hydrophobic barriers to this paper using a Xerox Colorcube, which allows to direct and control the flow in the paper device (Lee and Gomez, 2017; Lisowski and Zarzycki, 2013). In this case, we can push the liquid through the filter layer and to limit the loss of the sample through adsorption into each layer.

**Fig. 5.**
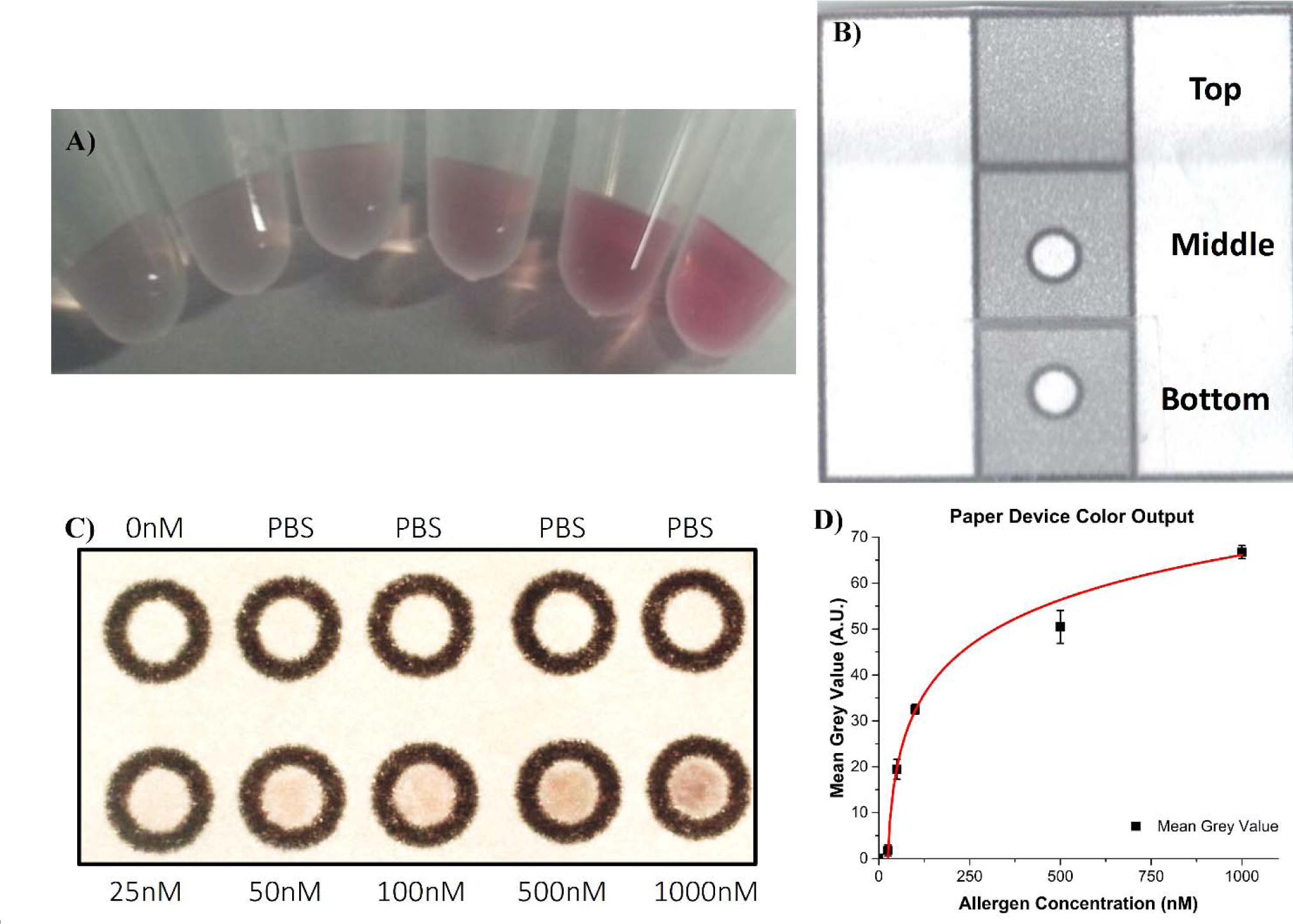
A) Image of the supernatant produced after simple filtration with Whatman 1 filter paper of samples with varying quantities of allergenic protein and the addition of GO, from left to right: 0 nM, 25 nM, 50 nM, 100 nM, 500 nM and 1000 nM of allergenic protein. B) Image of simple 3-fold paper based device for assistance in detection of allergens in food. The Top: Lid Layer, Middle: Filter Layer, Bottom: Output Layer. C) Image of the output layer after filtering samples with varying amounts of allergenic proteins, from left to right: 0 nM, 25 nM, 50 nM, 100 nM, 500 nM and 1000 nM of allergenic protein. D) Graph of the measured mean grey value of ROI from image C.

For our design, we were able to utilize less than 30 μL of the GO reacted systems to view a measurable result. A scaled-up model with a greater number of testing zones was used to simultaneously test many allergen concentrations and peanut aptamer functionalized AuNP. We used various peanut allergen concentration samples and incubated 20 μL of the functionalized AuNP and then added GO. After 30 minutes the 20 μL of solution, which contained both the floating complexes and unbound AuNP that have developed, were dispersed on a layer of paper with designated zone made by printed hydrophobic wax. This layer is where the sample is first dispersed and acts as the filtering layer for the complexes, identical to the function of the middle layer in Fig. 5B. This layer was then placed on top of another layer, of identical design, with hydrophobic barrier defined zones. The backside of this bottom layer was covered with tape to prevent any further fluid transfer. As shown in Fig. 5B, this bottom layer is designed to function as the output readout or the color presentation layer. The last top cover layer, which was simply a sheet of para film wax was placed on the top. This acts similarly to the top layer in Fig. 5B and is used to help push the sample through the filter layer onto the color output readout layer. A large sheet of glass placed on the top to help to provide even pressure to all testing areas as we pushed by hand. After 1 minute, the layers were disassembled and the output layer, as seen in Fig. 5C, shows the resultant dying by the free AuNP. Visibly, this has made direct observation much simpler and can help the user quickly identify the possible level of contamination. We can further quantify the results using very simple processing procedures, as conducted by (Jokerst et al., 2012). Requiring only an image of the testing zone to measure average mean grey intensity. Using ImageJ software (NIH, USA), we collected information on the mean grey average of each zone after testing. Fig. 5D is the collection of this data, showing us a similar output to that of the in-solution verification (Fig. 4B). From this we can also quantify lower levels of allergen concentration with significant reliability, though the requirement of a camera is necessary. Although this is the case, the availability and accessibility of smart devices with a camera in comparison to a bench top spectra device is easily seen. The simple processing steps to acquire the mean grey average from the output zone in an image is far simpler and more accessible. Using the method described by Thomsen et al., (2003) we calculated the lowest posssible detectable concentation with a 3σ signal to noise ratio for the paper device, being found to be 27 nM. Although we can detect down to 25 nM using the spectra, the paper devices showed reproducible identification for values at or above 50 nM with the peanut aptamer functionalized AuNP. Below this amount it became difficult to see or measure any color dying caused by AuNP which were not absorbed into a complex with GO in the paper based device.

## 4. Conclusions

For individuals living with allergen sensitivity, there is a significant lack of pro-active measures to help protect their health and well-being. There is a necessity for a bio-sensor system that can directly benefit both government and manufacturers, that is accurate, specific, low-cost and requires limited usage of laboratory instrumentation to detect allergen contamination in food. By utilizing the power of golds strong optical properties, with the selectivity and sensitivity of aptamer and the power of paper-based microfluidics the proposed device can meet these needs. With the proposed mechanism, we could accurately measure allergens from 25 nM - 1000 nM with LoD of 7.8 nM, 12.4 nM and 6.2 nM for peanut, milk and shrimp allergens respectively in solution. A 50 nM - 1000 nM range with a LoD of 27 nM was achieved when integrated with the microfluidic paper device when utilized with a smart device with a camera. Furthermore, the system only requires the need for two reagents, functionalized AuNP and GO, and a simple and cheap paper based device. Further optimization and investigation is however required to both fully understand the capabilities of the mechanism and to create a better experience for the end user. Overall, the proposed device is highly comparable in time to other quantitative methods, such as ELISA, with even greater sensitivity and less complexity. This combination allows it to be a potential tool for those living with allergen sensitivities and giving food manufacturer’s an in-expensive and efficient method to control cross-contamination and mitigate associated economic losses.

## Acknowledgments

The authors sincerely thank the Natural Sciences and Engineering Research Council of Canada (#400705) for funding this study.

